# PTRN-1/CAMSAP and NOCA-2/NINEIN are required for microtubule polarity in *Caenorhabditis elegans* dendrites

**DOI:** 10.1101/2021.12.13.472373

**Authors:** Liu He, Lotte van Beem, Casper C. Hoogenraad, Martin Harterink

## Abstract

The neuronal microtubule cytoskeleton is key to establish axon-dendrite polarity. Dendrites are characterized by the presence of minus-end out microtubules, however the mechanisms that organize these microtubules minus-end out is still poorly understood. Here, we characterized the role of two microtubule minus-end related proteins in this process in *Caenorhabditis elegans*, the microtubule minus-end stabilizing protein CAMSAP (PTRN-1) and a NINEIN homologue (NOCA-2). We found that CAMSAP and NINEIN function in parallel to mediate microtubule organization in dendrites. During dendrite outgrowth, RAB-11 positive vesicles localized to the dendrite tip function as a microtubule organizing center (MTOC) to nucleate microtubules. In the absence of either CAMSAP or NINEIN, we observed a low penetrance MTOC vesicles mis-localization to the cell body, and a nearly fully penetrant phenotype in double mutant animals. This suggests that both proteins are important for localizing the MTOC vesicles to the growing dendrite tip to organize microtubules minus-end out. Whereas NINEIN localizes to the MTOC vesicles where it is important for the recruitment of the microtubule nucleator γ-tubulin, CAMSAP localizes around the MTOC vesicles and is co-translocated forward with the MTOC vesicles upon dendritic growth. Together, these results indicate that microtubule nucleation from the MTOC vesicles and microtubule stabilization are both important to localize the MTOC vesicles distally to organize dendritic microtubules minus-end out.

## Introduction

In neurons, the microtubule cytoskeleton is vital for axon and dendrite development. In axons microtubules are mainly arranged with their plus-ends distal to the cell body, whereas dendritic microtubules are predominantly arranged with their minus-ends distal to the cell body in invertebrates or have mixed orientation microtubules in vertebrates (Baas et al., 1988; Kapitein and Hoogenraad, 2015; Stepanova et al., 2003; Stone et al., 2008). This difference in microtubule organization allows for selective cargo transport into axons or dendrites (Rolls and Jegla, 2015). Defects in this organization lead to protein trafficking defects and to defects in neuron development and functioning (Harterink et al., 2018; He et al., 2020; Yogev et al., 2016). Although the importance of the microtubule cytoskeleton organization is apparent, the molecular mechanism controlling differential microtubule organization between axons and dendrites is still not fully clear.

During cell division, the centrosome is the main microtubule organizing center (MTOC). However, in polarized cells such as neurons, most microtubules are organized in a non-centrosomal manner (Nguyen et al., 2011). Several mechanisms have been proposed to organize the neuronal microtubule cytoskeleton. These include the transport of microtubules into the correct organization (also referred to as microtubule sliding), the local nucleation of microtubules and the selective stabilization of correctly organized microtubules by microtubule associated proteins (MAPs) (Baas and Lin, 2011; Baas et al., 2006; Baas and Yu, 1996; Fréal et al., 2019; He et al., 2020; Kapitein and Hoogenraad, 2015; Liang et al., 2020; Maniar et al., 2011; Weiner et al., 2020). The formation of the axon is typically regarded as the initial step in neuron polarization. Stabilization of axonal microtubules is a critical early event to form the axon. Indeed, the artificial stabilization of microtubule using drugs leads to the induction of multiple axons (Gomis-Rüth et al., 2008; Witte et al., 2008) and the formation of axons in mammals relies amongst others on the TRIM46 protein that stabilizes and bundles the microtubule in a parallel plus-end out fashion in the proximal axon (Fréal et al., 2019; Harterink et al., 2019; van Beuningen et al., 2015). This plus-end out microtubule organization can be propagated into the growing axon by several mechanisms: by the outgrowth of existing microtubules potentially followed by severing (Ahmad et al., 1999; Kapitein and Hoogenraad, 2015); by forward translocation of the microtubule bundle (Roossien et al., 2014) and by the local nucleation of new microtubules along the lattice of pre-existing microtubules using the Augmin complex (Cunha-Ferreira et al., 2018; Sánchez-Huertas et al., 2016). The transport of small microtubule fragments from the cell body has also been observed but so far it is not known if this contributes to the propagation of the axon cytoskeleton (Hasaka et al., 2004; Wang and Brown, 2002).

How dendrites acquire and maintain their typical minus-end out oriented microtubules is less clear; although Augmin mediated microtubule nucleation and various MAPs were shown to contribute to the mixed microtubule organization in mammalian neurons (Cunha-Ferreira et al., 2018; Kapitein and Hoogenraad, 2015; Maniar et al., 2011; Sánchez-Huertas et al., 2016; Yau et al., 2014). In invertebrate neurons, the uniform minus-end out microtubule organization may offer a simpler starting point to understand the origin of the minus-end out microtubules. Early work suggested a role for microtubule sliding; in *Drosophila* microtubules can be slid by kinesin-1 during the earliest phases of neuron development (Winding et al., 2016a) and in *C. elegans* a mutant for the motor kinesin-1 loses minus-end out microtubules in dendrites (Yan et al., 2013). However, lately the role of local microtubule nucleation has gained attention, since non-centrosomal MTOCs are found localized in the dendrites and may be important to locally nucleate the minus-end out microtubules (Liang et al., 2020; Valenzuela et al., 2020; Weiner et al., 2020; Weiner et al., 2021). For example, Golgi outposts and early endosomes have been shown to nucleate microtubules in *Drosophila* dendrites and were suggested to contribute to the minus-end out microtubule organization in these neurites (Mukherjee et al., 2020; Ori-McKenney et al., 2012; Weiner et al., 2020; Ye et al., 2007). Shortly thereafter, in *C. elegans*, it was shown that RAB-11 positive endosomes localize to the dendrite growth cone to nucleate microtubules with minus-end out organization (Liang et al., 2020). It is unclear however, how the microtubule nucleating γ-tubulin is recruited to the RAB-11 vesicles and how the minus-end out microtubules are specifically maintained in dendrites.

A prominent protein family regulating microtubule stabilization is the CAMSAP family: CAMSAP1-CAMSAP3 (in vertebrates), Patronin (in *Drosophila*), and PTRN-1 (in *C. elegans*). These proteins can bind microtubule minus-ends and thereby protect them against depolymerization (Akhmanova and Hoogenraad, 2015; Goodwin and Vale, 2010; Jiang et al., 2014; Meng et al., 2008). Indeed, in cultured mammalian neurons, microtubule stabilization by CAMSAP proteins was found critical for neuronal polarization (Pongrakhananon et al., 2018; Yau et al., 2014; Zhou et al., 2020) and also in *Drosophila* Patronin was found important for dendritic microtubule polarity (Feng et al., 2019). In addition to CAMSAP, microtubules can be stabilized by microtubule associated proteins (MAPs) that can crosslink microtubules together or connect them to cortical structures via Ankyrin or Spectraplakin proteins (Fréal et al., 2019; Prokop et al., 2013). For example, we found that in *C. elegans* the UNC-33(CRMP)/UNC-119/UNC-44(Ankyrin) complex connects the microtubule cytoskeleton to the cortex in both axons and dendrites to maintain the proper polarity organization (He et al., 2020).

In this study we found that PTRN-1 (CAMSAP) is important in *C. elegans* for the proper localization of the MTOCs vesicles and therefore minus-end out microtubule polarity in the growing dendrite. Moreover we found that the Ninein homologue NOCA-2 acts in parallel to localize γ-tubulin to the MTOC vesicles. Our results suggest that microtubule nucleation from MTOCs vesicles acts together with microtubule stabilization by CAMSAP proteins to organize dendritic microtubules minus-end out.

## Results

### Microtubule organization during early neuronal development depends on PTRN-1 (CAMSAP) and UNC-33 (CRMP)

The *C. elegans* PVD neuron is an excellent model to study neuron development in vivo (Sundararajan et al., 2019). The microtubule cytoskeleton is for a large part restricted to the primary branches where it is organized with minus-end out polarity in the anterior dendrite and with plus-end out polarity in the axon and posterior dendrite (Figure 1A) (Harterink et al., 2018; Taylor et al., 2015a). To address how these differences in microtubule organization are set up and maintained, we compared the contribution of two microtubule stabilizing proteins using genetic mutants; UNC-33 (CRMP) and PTRN-1 (CAMSAP). To visualize microtubule orientation, we used the microtubule plus-end protein EBP-2 fused to GFP or mKate2, which localizes to the microtubule plus-end only during microtubules growth. We found that both *unc-33* and *ptrn-1* mutations affect the microtubule organization in the anterior dendrite of the mature PVD neuron but in a distinct manner (Figure 1B-D). Whereas loss of *unc-33* led to a fully penetrant mixed or reversed microtubule polarity phenotype (as reported before (He et al., 2020; Maniar et al., 2011)), the loss of *ptrn-1* led to occasional full reversal of microtubule polarity (Figure 1B-1D). This suggests that these proteins function differently in organizing neuronal microtubules. In the non-ciliated PHC neuron or the ciliated URX neuron we did not observe microtubule organization defects in the *ptrn-1* mutant (Supplemental figure 1A-B), which suggests that these neurons do less or do not dependent on PTRN-1.

**Figure 1.**
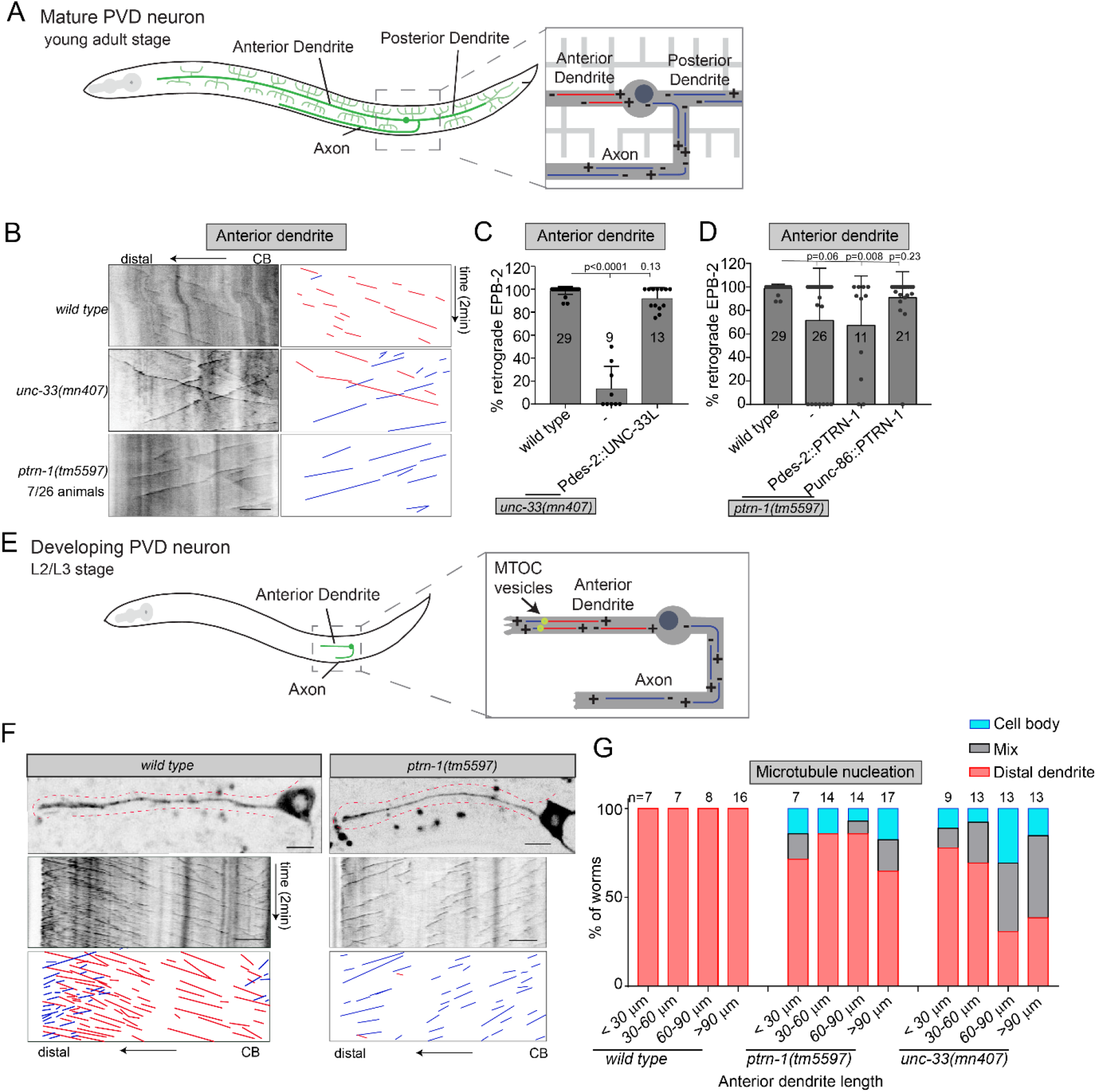
Initial microtubule polarity establishment partially depends on PTRN-1 (CAMSAP) and UNC-33 (CMRP) (A) Schematic representation of the microtubule organization in mature PVD neuron; the anterior dendrite contains uniform minus-end out microtubules (red), whereas the axon and posterior dendrites contain uniform plus-end out microtubules (blue)(Harterink et al., 2018). (B-D) Analysis of microtubule organization in the proximal PVD anterior dendrite using EBP-2::GFP to visualize the microtubule plus-end growth events. (B) Kymograph illustrating the EBP-2::GFP growth events in the PVD anterior dendrite; 7 out of 26 animals have completely reversed microtubules orientation in *ptrn-1* mutant (C) Quantification of the percentage of retrograde growth evens in *unc-33* mutant with or without the *des-2* promoter driven rescue construct (late expression). (D) Quantification of the percentage of retrograde growth evens in *ptrn-1* mutant with or without the *des-2* (late expression) or *unc-86* promoter (early expression) driven rescue constructs. Scale, 5 μm. Analyzed animals were from the L4 or young adult stage; error bars represent SD; statistical analysis, Kruskal-Wallis test followed by Dunn’s multiple comparisons test. Number of analyzed animals is indicated. (E) Schematic representation of the microtubule organization in the developing PVD neuron. In the anterior dendrite, a distal MTOC localized in the anterior dendrite generates dendritic minus-end out microtubules and short plus-end out microtubules that grow towards the dendritic tip (Liang et al., 2020). (F-G) Analysis of microtubule organization in the developing PVD anterior dendrite using EBP-2::GFP. Representative kymographs are shown (F) and the main site of microtubule growth events was quantified as the neuron develops (G). Scale, 5 μm; the number of analyzed animals is indicated; red line indicates the growing dendrite.

We previously found that the *unc-33* (CRMP) mutant phenotype can be rescued by reexpression of UNC-33L using the *des-2* promoter (Figure 1C) (He et al., 2020). As this promoter drives expression after the initial neurite outgrowth (Maniar et al., 2011), this suggests that UNC-33 is important at later stages of neuronal development. In contrast, we found that the *ptrn-1* (CAMSAP) mutant phenotype cannot be rescued by reexpression of PTRN-1 using the late *des-2* promoter but it can be rescued using the early expressing *unc-86* promoter (Figure 1D) (Baumeister et al., 1996). This suggests that PTRN-1 functions earlier than UNC-33 to organize microtubules in the PVD neuron.

It was recently shown that during the outgrowth of the anterior PVD dendrite, distal RAB-11 positive vesicles function as a microtubule organizing center (MTOC) to nucleate microtubules towards the cell body to organize the dendritic microtubules minus-end out (Figure 1E)(Liang et al., 2020). Indeed when imaging EBP-2::GFP dynamics during early dendrite development we saw pronounced microtubule growth events in the distal dendrite in both anterograde and retrograde directions (Figure 1F). Interestingly, we observed similar distal microtubule nucleation in most *unc-33* mutant animals (Figure 1G), even though the adult animals have a fully penetrant phenotype (Figure 1C). We observed a gradual increase in the phenotype upon neuron maturation (Figure 1 G), suggesting that UNC-33 is continuously required to organize the microtubules. Since UNC-33 (CRMP) forms a complex with UNC-44 (Ankyrin) and UNC-119 (UNC119) to anchor microtubules to the cortex (He et al., 2020), and that we found similar defects in the *unc-119* mutant (Supplemental figure 1D), this suggests that loss of cortical microtubule anchoring is the cause of this increasing phenotype. In the *ptrn-1* (CAMSAP) mutant a small population of animals lost the pronounced distal microtubule growth events at early developmental stages and had reversed microtubule polarity organization (Figure 1F-G). In contrast to *unc-33* mutants, the fraction of animals with this defect remained constant during development and is similar to the defects seen in adult animals (Figure 1D). This suggest that PTRN-1 mainly acts at early stages to ensure distal microtubule nucleation to set up minus-end out microtubule polarity.

Together, these results indicate that the microtubules stabilizing proteins UNC-33 (CRMP) and PTRN-1 (CAMSAP) both act in the PVD neuron to regulate microtubules organization. However, they function differently: PTRN-1 mainly functions at early stages whereas UNC-33 acts continuously during neuron development to maintain microtubule polarity by connecting the microtubule cytoskeleton to the cortex.

### ptrn-1 (CAMSAP) and noca-2 (NINEIN) function in parallel to organize microtubules minus-end out in dendrites

Since loss of *ptrn-1* (CAMSAP) only shows a partially penetrant phenotype, this suggests that other factors work in parallel to set up dendritic microtubule organization in the PVD neuron. Previously, *ptrn-1* was shown to act in parallel to *noca-1* (NINEIN) to organize the non-centrosomal microtubule organization in the *C. elegans* epidermis (Wang et al., 2015). NOCA-1 has some sequence and functional similarity to mammalian NINEIN (Wang et al., 2015). BLAST searches using mammalian NINEIN however, identify another protein in *C. elegans*, which we called NOCA-2. Using the sensitive homology prediction software HHpred (https://toolkit.tuebingen.mpg.de/tools/hhpred), the NOCA-2 homology resides in the N-terminal domain (~500 amino acids) of mammalian NINEIN and NINEIN-like protein. Using CRISPR we generated the *noca-2(hrt28*) mutant, which has a 4782bps deletion encompassing the entire coding sequence (Supplemental figure 1C). We found that single mutants for either *noca-1* or *noca-2* showed a low penetrance microtubule polarity reversal phenotype in the mature PVD neuron, similar to the *ptrn-1* mutant (Supplemental figure 2A-B and Figure 2A-B). This defect seems specific to the minus-end out microtubules, as the microtubule polarity in the PVD axon and posterior dendrites were not affected (Supplemental figure 2C-F). Combining *prtn-1* with either *noca-1* or *noca-2*, led to a strong enhancement of the phenotype (Figure 2A-B and Supplemental figure 2A-B). This suggest that *ptrn-1* acts in parallel to *noca-1* and *noca-2* in the PVD neuron, similar to what was observed before in the epidermis for *ptrn-1* and *noca-1* (Wang et al., 2015). The *ptrn-1;noca-2* double mutant animals superficially look like wildtype animals, however the *ptrn-1;noca-1* double mutant worms exhibit severe developmental defects (Supplemental figure 2H) (Wang et al., 2015). Therefore, we cannot rule out that the neuronal defects are secondary to defects in the surrounding epidermis. Moreover, we were not able to rescue the *noca-1* mutant phenotype using two previously generated functional tagged transgenes (Supplemental figure 2G) (Wang et al., 2015), therefore we chose to here focus our studies on *noca-2.* Reexpression of NOCA-2 in the PVD neuron using the early expressing *unc-86* promoter was able to fully suppress the microtubule defect (Figure 2C), which may suggest that *noca-2* is also needed during early PVD development for proper microtubule organization. Indeed, looking at early dendrite development, we observed a low percentage of *noca-2* mutant animals that completely lost the distal microtubule dynamics (Figure 2E). This phenotype that was strongly enhanced in the *ptrn-1;noca-2* double mutant (Figure 2D-E, Video 1), similar to the mature neuron situation (Figure 2B). Quantification of the distal microtubule dynamics using EBP-2::GFP or GFP::TBA-1 (Tubulin) in the animals that retained the distal dynamics, show reduced growth events mainly in the *noca-2* mutant background (Supplemental figure 3A-D, movie1). Similarly, the microtubule plus-end growth speed and distance decreased upon *noca-2* depletion, whereas *ptrn-1* only had no or very subtle defects (Supplemental figure 3A-B, Video 1). This suggest that although the *noca-2* and *ptrn-1* single mutants have similar microtubule organization defects in the mature neuron, these proteins act differently during development.

**Figure 2.**
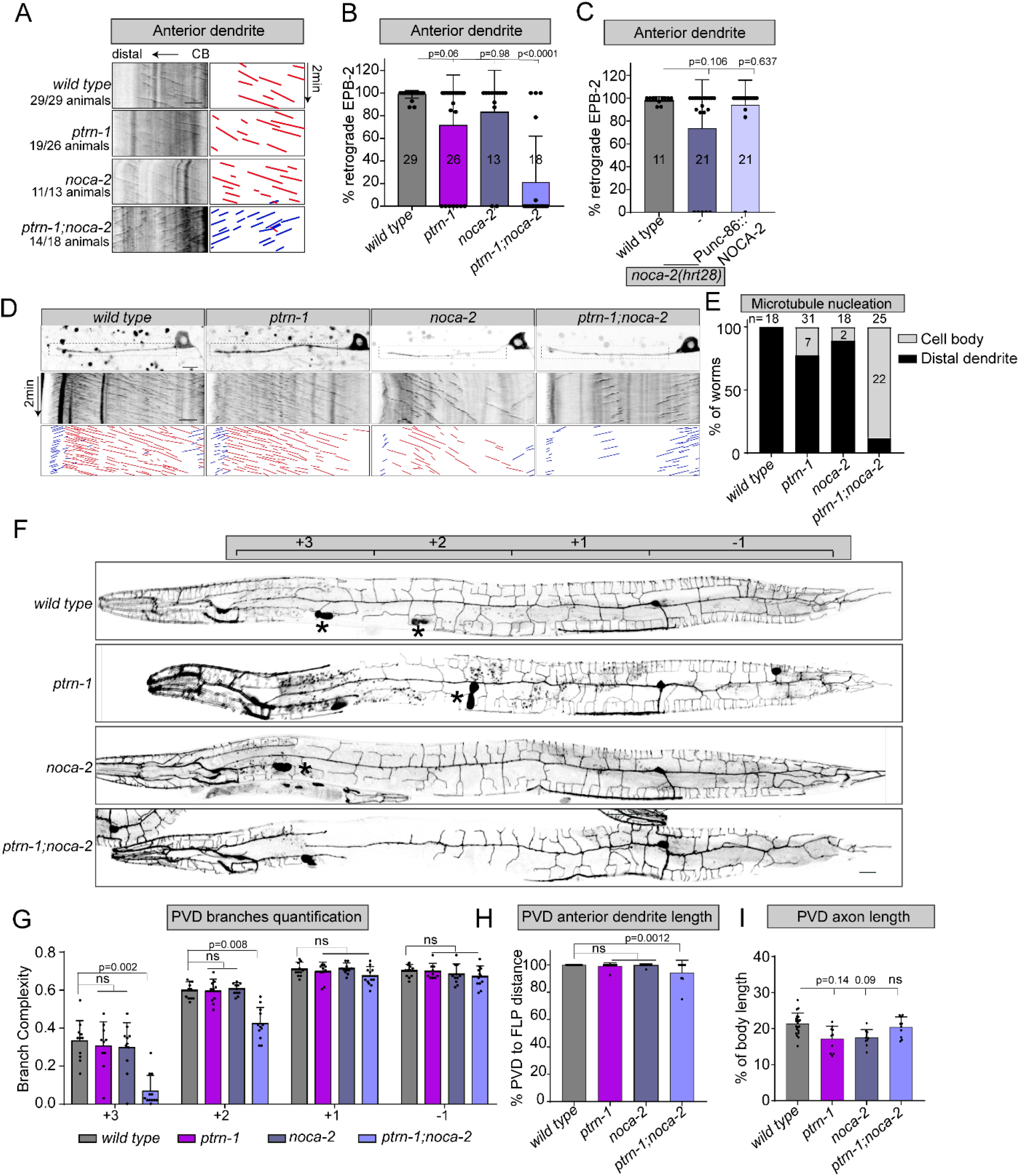
PTRN-1 (CAMSAP) and NOCA −2 (NINEIN) act in parallel to organize microtubule minus-end out in dendrites. (A-C) Analysis of microtubule organization in the mature PVD anterior dendrite using EBP-2::GFP to visualize the microtubule plus-end growth events. (A) Kymographs illustrating EBP-2::GFP growth events in the PVD anterior dendrite in the wild type and indicated mutants. The numbers of animals about this phenotype is indicated. (B) Quantification of the retrograde growth events in mature PVD anterior dendrite in the wild type and indicated mutant backgrounds. (C) Quantification of the percentage of EBP-2 retrograde growth evens in *noca-2* mutant with or without the *unc-86* promoter (early expression) driven rescue construct. Scale, 5 μm. Analyzed animals were from the L4 or young adult stage; error bars represent SD; statistical analysis, Kruskal-Wallis test followed by Dunn’s multiple comparisons test. Number of analyzed animals is indicated. (D-E) Analysis of microtubule organization in the developing PVD anterior dendrite using EBP-2::GFP in indicated mutants. Representative kymographs are shown (D) and the main site of new growth events was quantified (E). Scale, 5 μm. Error bars represent SD; statistical analysis, Kruskal-Wallis test followed by Dunn’s multiple comparisons test. Number of analyzed animals is indicated. (F-I) quantification of PVD morphology in the indicated mutants. (F) Representative images of the PVD morphologies. Note that this marker is also expressed in another neuron (FLP) located in the head (left of the +3 region) and also in the coelomocytes (marked with *) which were used as injection marker. (G) Quantification of PVD dendritic branch complexity in the four PVD regions along the anteroposterior axis as indicated in (F), based on (Caitlin A. Taylor et al, 2015, plos genetic). (H) Quantification of the anterior dendrite outgrowth towards the FLP cell body localized in the head. (I) Quantification of the relative axon length in the ventral nerve cord. Scale, 20 μm. Analyzed animals were young adult stage; Error bars represent SD; statistical analysis, Kruskal-Wallis test followed by Dunn’s multiple

Since the microtubule organization is critically important for development and growth of neurons (Barnes and Polleux, 2009; Kapitein and Hoogenraad, 2015), we analyzed the morphology of the mature PVD neuron. We observed that PVD neuron shows little defects in the *ptrn-1* or *noca-2* single mutant, while the *ptrn-1;noca-2* double mutant displayed pronounced morphological defects (Figure 2F); it had reduced dendritic complexity especially in the anterior segments (Figure 2G) and the primary anterior dendrite was shorter (Figure 2H). We did not observe obvious defects in the posterior dendrite or the axon (Figure 2G and 2I).

Taken together, these results show that PTRN-1 (CAMSAP) and NOCA-2 (NINEIN) act in parallel in the PVD neuron during early development to establish minus-end out microtubule organization, and that this organization is important for proper dendritic morphogenesis.

### PTRN-1 and NOCA-2 are essential for MTOC vesicles transport during dendrite outgrowth

It was previously shown that the microtubule nucleator γ-tubulin is transported on RAB-11 positive endosomes to the tip of the developing dendrite in the PVD neuron to establish the minus-end out microtubules (Liang et al., 2020). Indeed, we detected dynamic GIP-2::GFP puncta (γ-tubulin small complex subunit*2* (Wang et al., 2015)) in the distal segment of the anterior dendrite (Figure 3A and Supplemental figure 5A) and these puncta perfectly overlapped with the RAB-11 marker (Figure 3B-C). To further investigate whether the microtubule polarity defects we observed in the *ptrn-1* (CAMSAP) and *noca-2* (NINEIN) mutants were caused by mislocalization of RAB-11 endosomes, we performed time-lapse imaging of PVD expressed RAB-11::GFP during neuron development. In the wild type animals, RAB-11 vesicles were consistently presents at the tip of the growing dendrite (Figure 3D and 3F, Video 2). In the *ptrn-1* mutant, some animals lost the distal RAB-11 vesicles although most animals (13 out of 15) kept RAB-11 vesicles localized in the growing anterior dendrite (Figure 3D and 3F). Similarly in absence of NOCA-2, 24 out of 29 animals have RAB-11 vesicles distally localized in anterior dendrite (Figure 3D and 3F, Video 2). In contrast to the single mutants, in the double mutant RAB-11 vesicles were almost entirely lost from the distal dendrite and instead localize to the cell body (Figure 3D and 3F, Video 2). These defects closely reflect the microtubule defects we have observed in the mutants and strongly suggests that the loss of minus-end out microtubules in the dendrites of *ptrn-1* and *noca-2* mutants is caused by a loss of RAB-11 localization to the distal dendrite during outgrowth.

**Figure 3.**
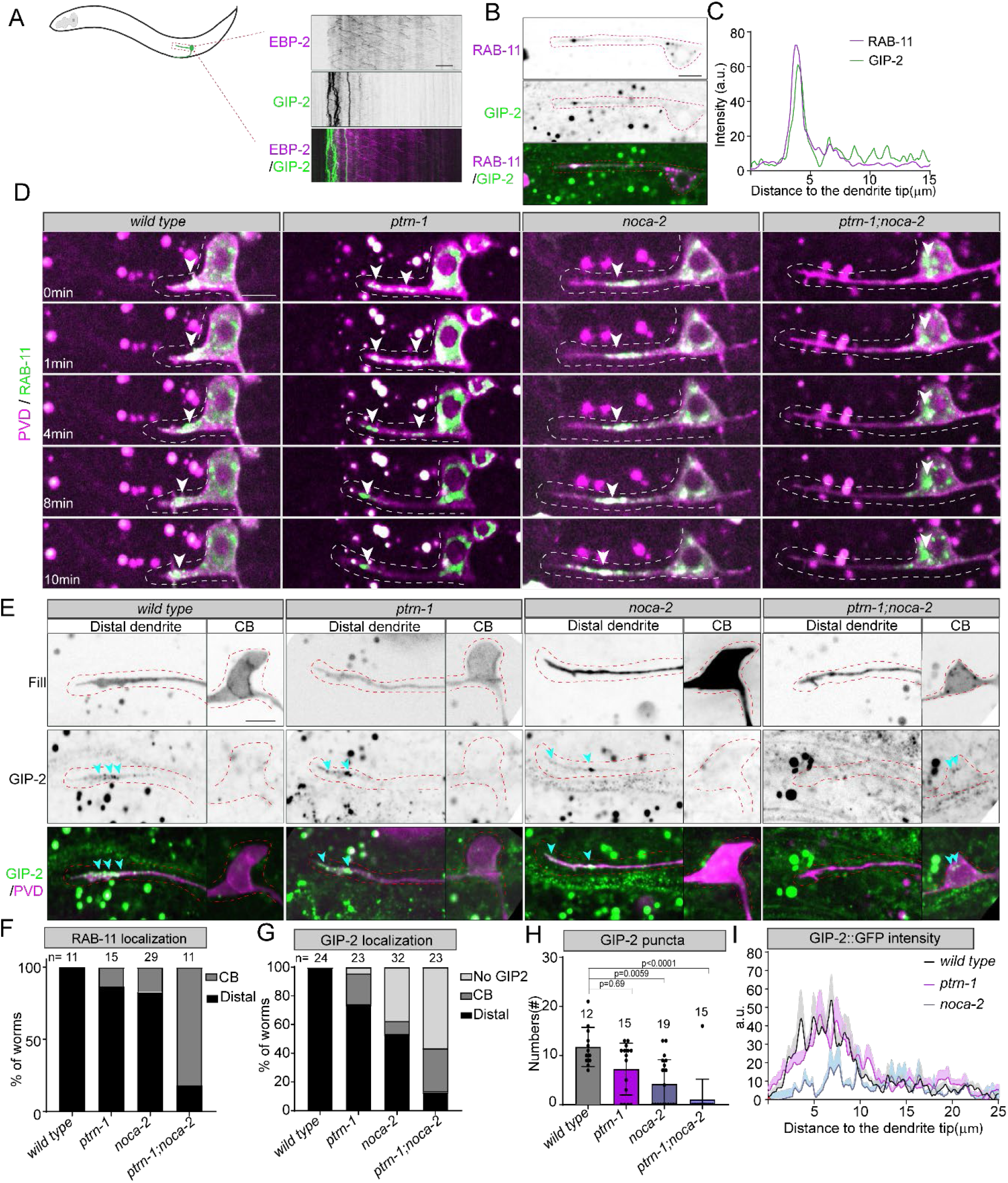
PTRN-1 and NOCA-2 are essential for distal MTOC vesicle localization during neurites outgrowth. (A) Kymographs of the growing PVD anterior dendrite expressing EBP-2::mKate2 (PVD specific) and GIP-2::GFP (knock-in). Scale, 5 μm. (B-C) Co-localization of GIP-2::GFP (knock-in) and mKate-2::RAB-11 (PVD expressed) in distal segment of the growing PVD anterior dendrite. And the line scans for intensity profile of each channel from distal dendrites to cell body is showed in(C). Scale, 5 μm. PVD neurons are indicated with red dashed lines in(B). (D) Vesicle dynamics of PVD expressed GFP::RAB-11 in distal segment of the growing anterior dendrite in wild type and indicated mutants. White arrowheads point to the dynamic RAB-11 clusters. Scale, 5 μm. Dashed line marks the developing dendrite. (E) GIP-2::GFP (knock-in) localization in the wild type and indicated mutants; green: GIP-2,magenta: fill of PVD neurons. GIP-2 cluster are indicated with blue arrowheads and the PVD neurons are indicated with red dashed lines. Scale, 5 μm. (F-G) Quantification of RAB-11 (D) and GIP-2 (E) localization in wild type and indicated mutants in the developing PVD neuron; light gray: no GIP-2 cluster was observed, dark gray: RAB-11 or GIP2 cluster localized in cell body, black: RAB-11 or GIP-2 accumulated in the distal developing dendrites. Number of analyzed animals is indicated. (H-I) Quantification of number of GIP-2::GFP puncta (H)) and the line scans of average intensity from dendritic tip to cell body in distal segment of the growing PVD anterior dendrite (I).Error bars represent SD; statistical analysis, Kruskal-Wallis test followed by Dunn’s multiple comparisons test. Number of analyzed animals is indicated.

In parallel we also analyzed γ-tubulin localization using a GIP-2::GFP knock-in line (Wang et al., 2015). As it is expressed in all tissue and for a large part diffusely localized in the cells, only places with clear enrichment can be detected above the background. In wildtype animals we consistently observed GIP-2 puncta in the distal dendrite (24 animals). In the *ptrn-1* (CAMSAP) mutant, GIP-2 puncta mis-localized to the cell body in 5 out of 23 animals (Figure 3E and 3G, Supplemental figure 4A), which is consistent with the RAB-11 localization defects (Figure 3F) and microtubule polarity defect (Figure 1D and 1G). In the *noca-2* (NINEIN) mutant however, in 12 out of 32 animals we could not detect obvious GIP-2 clusters neither at the dendrite tip nor in the cell body (Figure 3G and Supplemental figure 4A). This suggests that γ-tubulin fails to be efficiently recruited to RAB-11 vesicles to sufficient levels that can be detected above background levels. When analyzing RAB-11 co-localization with GIP-2 in the *ptrn-1* and *noca-2* mutants, we indeed found that in the *noca-2* mutant 5 out of 17 animals had a distal RAB-11 cluster without obvious GIP-2 enrichment, whereas in the *ptrn-1* mutant all distal RAB-11 vesicles co-localized with GIP-2 (Supplemental figure 4B-C). Not surprisingly, *noca-2* mutants have a reduced number of GIP-2 puncta in the distal dendrite (Figure 3H) and reduced overall GIP-2 levels (Figure 3I), which is in agreement with the fact that specifically in *noca-2* mutants we observed reduced microtubules dynamics in the distal dendrite (Supplemental figure 3A-D).

In conclusion, these results suggest that *noca-2* (NINEIN) works in parallel to *ptrn-1* (CAMSAP) to mediate distal MTOC localization to the growing dendrite but that the proteins act differently. Whereas CAMSAP proteins are well described microtubule minus-end binding and stabilizing proteins that in the PVD dendrite may stabilize tracks for MTOC vesicle transport, the function of NOCA-2 seems related to the efficient recruitment of γ-tubulin to the MTOC vesicles.

### NOCA-2 (NINEIN) acts at the MTOC vesicles

In order to localize NOCA-2 (NINEIN) protein we used CRISPR-CAS9 to endogenously insert a GFP tag at the C-terminus of the NOCA-2 locus. We observed NOCA-2 accumulations in several spots in the head and tail of the animal (Figure 4A) and also at the cortex of the epidermal seam cells (Figure 4B). Indeed, using the NOCA-2 promoter to drive mKate2 we observed a broad expression (Supplemental figure 4D). Importantly, we detected NOCA-2::GFP accumulations in the distal PVD dendrite during neuron development, which showed a similar localization pattern and dynamics as GIP-2 and RAB-11 (Figure 4C-D, Video 3). In order to validate that NOCA-2 localizes to the MTOC vesicles, we expressed RAB-11::mKate2 in the PVD and observed clear colocalization (Figure 4E-F). Since in the *noca-2* mutant γ-tubulin recruitment to the MTOC vesicles is reduced, this suggests that NOCA-2 localizes to the MTOC vesicles to recruit γ-tubulin for proper microtubule nucleation to organize dendritic microtubules minus-end out. How NOCA-2 in involved in recruiting γ-tubulin is so far unclear. Especially since in seam cells, we found that endogenous NOCA-2 only partially colocalizes with γ-tubulin (GIP-1) (Supplemental figure 4E-F) and that the γ-tubulin localization is not obviously changed in the *noca-2* mutant (Supplemental figure 4H). Moreover, the GIP-2 localization in seam cells was not affected when NOCA-2 was artificially localized to the mitochondria membrane (Supplemental figure 4G). This argues against a direct interaction to recruit γ-tubulin to the MTOC vesicles or that the interaction is regulated in a tissue specific manner.

**Figure 4.**
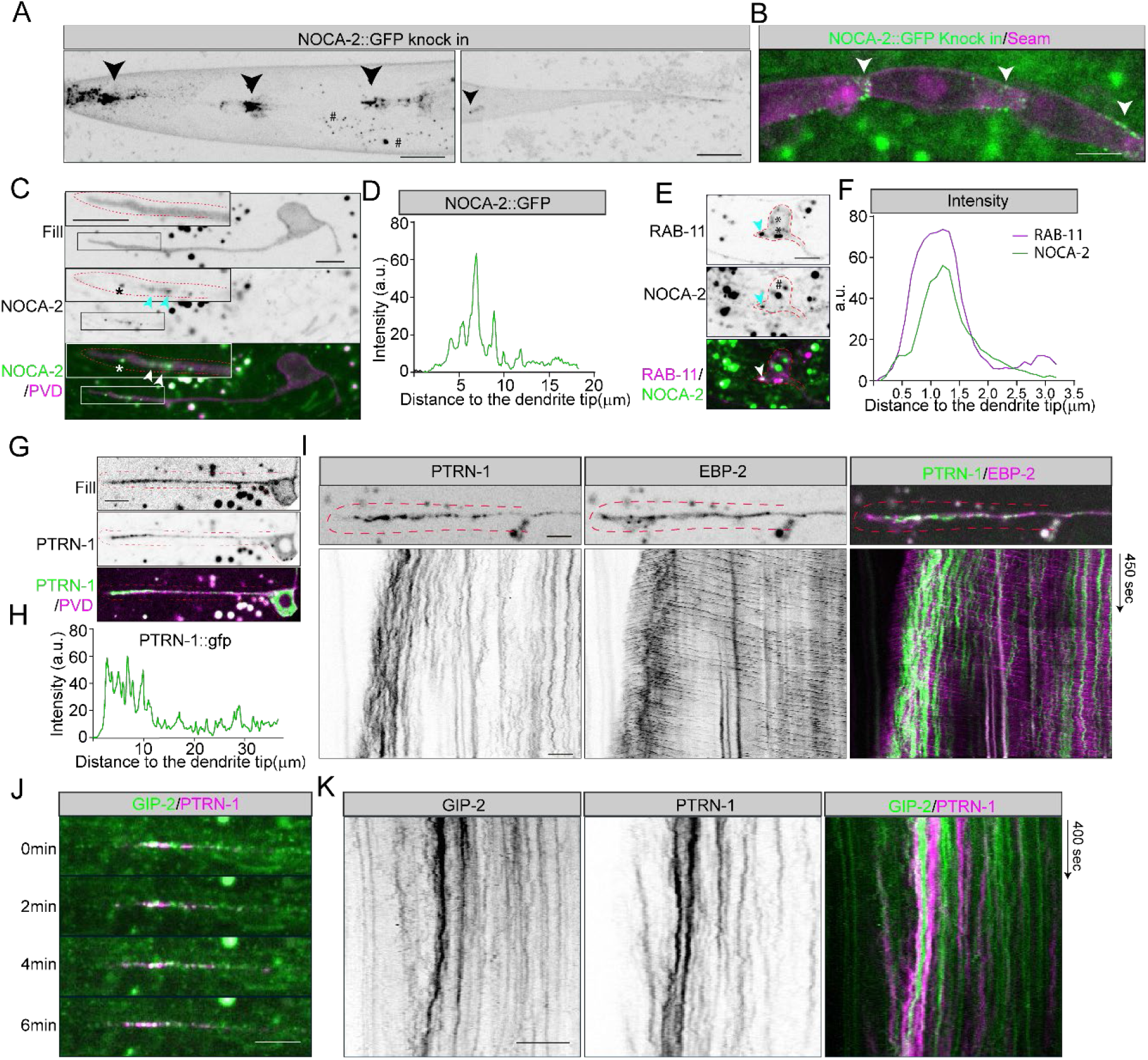
NOCA-2 localizes to the distal MTOC vesicles. (A-B) The localization of endogenously tagged NOCA-2::GFP. NOCA-2 accumulates to head and tail structures (marked with arrowheads) (A) and to junctions of epidermal seam cells(indicated with arrowheads)(B) (the puncta marked with # are autofluorescent gut-granules). Scale, 10 μm. (C-D) Endogenous NOCA-2::GFP localization (C) and the intensity quantification(D) in the PVD neuron during anterior dendrite outgrowth (magenta: neuron fill). NOCA-2 puncta are indicated with arrowheads in (C) and zoom in the top left (the puncta marked with * might be autofluorescent gut-granules). Scale, 5 μm. The developing anterior PVD dendrites are indicated with red dashed lines. (E-F) Co-localization (E) and the intensity profile (F) of endogenous NOCA-2::GFP (green) and PVD expressed mkate2::RAB-11 (magenta) in the developing PVD neuron. Arrowheads point to the co-localization of RAB-11 and NOCA-2. The puncta marked with # are autofluorescent and the RAB-11 puncta that without obvious NOCA-2 accumulation in PVD are marked with * in (E). Scale, 5 μm. The PVD neuron is indicated with red dashed lines. (G-H) Representative example (G) and intensity profile (H) of PVD expressed PTRN-1::GFP in the developing dendrite; Fill of PVD (magenta), PTRN-1 puncta (green). Scale, 5 μm. The anterior PVD dendrite is indicated with red dashed lines. (I) PTRN-1 dynamic in the growing PVD anterior dendrites with respect to microtubule nucleation; PTRN-1(green), EBP-2::GFP (magenta). Scale, 5 μm. The distal anterior PVD dendrites are indicated with red dashed lines. (J-K) co-transport and kymograph of GIP-2::GFP and PTRN-1::mKate2 in the distal segment of the growing PVD anterior dendrite. PTRN-1(magenta), GIP-2(green). Scale, 5 μm.

To further characterize the MTOC vesicles and better understand how NOCA-2 (NINEIN) and γ-tubulin may localize there, we have looked at two centrosomal proteins using GFP knock-in lines: the PCM scaffold protein SPD-5 and microtubule regulator TAC-1 (TACC) (Garbrecht et al., 2021; Magescas et al., 2021). However, neither of these proteins localizes to the MTOC vesicles in the PVD neuron (results not shown). This is consistent with a previous study that shows that SPD-5 mainly functions at the centrosomal MTOC (Kemp et al., 2004; Magescas et al., 2021; Magescas et al., 2019). Alternatively, NOCA-2 localization to the MTOC vesicles may be mediated by dynein. NINEIN proteins have been described as adaptors for dynein function (Casenghi et al., 2005; Redwine et al., 2017) and in the *C. elegans* PVD dynein was shown to localize to the MTOC vesicles to cluster these together (Liang et al., 2020). To investigate whether NOCA-2 functions with Dynein, we measured the width of GIP-2 cluster localized in the distal PVD dendrite in *noca-2* and *ptrn-1* (CAMSAP) mutants. We found that the GIP-2 cluster in the *noca-2* mutant is similar to the *ptrn-1* mutant, and only slightly wider than in wildtype animals (Supplemental figure 5B). In addition, distal DHC-1 (Dynein) accumulation didn’t show obvious difference in the *noca-2* mutant and the wild type (Supplemental figure 5C), arguing against a NOCA-2 link to Dynein. Therefore, more work is needed to understand how the RAB-11 positive vesicles in the PVD dendrite recruit the microtubule nucleation machinery.

### PTRN-1 puncta localize around the MTOC vesicles and translocate forward together

The involvement of the microtubule minus-end binding protein PTRN-1 (CAMSAP) in localizing the MTOC vesicles to the dendrite tip may suggest that PTRN-1 stabilizes a pool of microtubules in the PVD neuron that are important for the forward translocation of the MTOC vesicles. Interestingly, it was proposed that a pool of transiently stabilized plus-end out microtubules in the tip of the PVD dendrites serve as tracks for the kinesin-1 motor (UNC-116) to move the MTOC vesicles forward (Liang et al., 2020). To see if PTRN-1 could be involved in this process we expressed GFP or mKate2 tagged PTRN-1 in the PVD neuron. We observed that PTRN-1 puncta distributed throughout the dendrites with a clear enrichment in the distal dendrite around the MTOC vesicles with no obvious co-localization (Figure 4G-I and 4K). To see if MTOC vesicles may be transported over the PTRN-1 decorated microtubules we performed live imaging of PTRN-1. As expected, the PTNR-1 puncta in the mid-dendrite are largely immobile, suggesting that the microtubules are highly immobilized in the shaft (Figure 4I), e.g. by anchoring these to the cortex (He et al., 2020). We did see occasional (weak) anterograde moving PTRN-1 puncta (Supplemental figure 6D), which could represent minus-end growth (Feng et al., 2019; Puri et al., 2021). In the distal segment however the PTRN-1 puncta are more dynamic, suggesting a less anchored microtube cytoskeleton. More strikingly, when we imaged PTRN-1 together with the MTOC vesicle marker (GIP-2) we observed that their movement in the distal segments is correlated and that they translocate forward together upon dendrite growths (Figure 4J-K, Video 4). To see if the microtubule cytoskeleton is translocating forward in the distal dendrite, we analyzed a marker for stable microtubules (UNC-116(Rigor)::GFP) (Chen et al., 2021). The distal dendrite is largely devoid of the stable microtubule marker and only occasionally shows some microtubule dynamics (Supplemental figure 6A-B). This is consistent with the high EB and tubulin dynamics we observed in the distal segment (Figure 1F and Supplemental figure 3C-D) and suggests a mainly dynamic pool of microtubules. Using a marker for dynamic microtubules (UNC-104(Rigor)::GFP) over time proved to be hard, probably reflecting the high microtubule turnover (results not shown). Altogether these results argue against kinesin-1 transport over Camsap stabilized microtubules and instead suggests that the MTOC vesicles and the Camsap puncta are somehow connected. More work is needed to understand the mechanism that transports the MTOC vesicles forward.

## Discussion

The presences of minus-end out microtubules is one of the characteristic features that dendrites have over axons allowing for selective cargo transport to set up neuronal polarity. It was recently shown in *C. elegans* that microtubule nucleation from MTOC vesicles in the distal dendrite is essential to organize dendritic microtubules minus-end out (Liang et al., 2020). However how these vesicles localize to the growing dendrite tip is incompletely understood. Especially since these MTOC vesicles were suggested to be transported over the same microtubules as that they nucleate. This may suggest that these processes are connected and suggests a prominent role for selective microtubule stabilization. Here we report that the localization of these MTOC vesicles depends on the microtubule minus-end stabilizing protein PTRN-1 (CAMSAP), which functions in parallel to NOCA-2 (NINEIN) to set up dendritic minus-end out microtubules. Co-depletion of PTRN-1 with NOCA-2 leads to MTOC vesicles mislocalization to the cell body, which is the underlying cause of the microtubule polarity defects.

Since these proteins function (partially) redundantly, this suggests that these proteins act in different processes. Indeed, mutants for these genes have a different effect on microtubule properties in the distal dendrite and have a different localization pattern. PTRN-1 (CAMSAP) shows a punctate pattern throughout the dendrite, as expected for a protein that binds to microtubule minus-ends. We did see a clear enrichment of PTRN-1 puncta surrounding the MTOC vesicles suggesting that there is a higher microtubule density in the distal segment (Figure 4I). NOCA-2 (NINEIN) on the other hand localizes to the MTOC vesicles where it perfectly overlaps with the microtubule nucleator γ-tubulin (Figure 4E-F). Moreover, in the *noca-2* mutant we observed that γ-tubulin is not efficiently recruited to the MTOC vesicles (Figure 3G and Supplemental figure 4B-C), suggesting that NOCA-2 is involved in the recruitment of γ-tubulin for proper microtubule nucleation. However how NOCA-2 localizes to the vesicles and how it may recruit γ-tubulin is still an open question. Although the MTOC vesicles are RAB-11 positive, many RAB-11 positive endosomes localize to the cell body without obvious NOCA-2 accumulation (Figure 4E). Moreover RAB-11 depletion had only mild defects on GIP-2 localization (Liang et al., 2020), therefore, it seems unlikely that RAB-11 directly recruits NOCA-2 and suggests that other proteins are involved. The endosomal recruitment of NOCA-2 may be aided by palmitoylation, as was previously shown for NOCA-1 (Wang et al., 2015); http://lipid.biocuckoo.org/ predicts a high threshold palmitoylation site at C441 in NOCA-2. To recruit the γ-tubulin to the MTOC vesicle membrane, NOCA-2 may directly or indirectly interact with γ-tubulin, since pull down experiment in mammalian cells showed that NINEIN is in a complex with γ-tubulin (Delgehyr et al., 2005). However, the NOCA-2 to human NINEIN sequence conservation is low, therefore this will have to be tested for the *C. elegans* proteins as well. Moreover, we did not observe clear co-localization of γ-tubulin and NOCA-2 in other tissues (Supplemental figure 4E-G), arguing against a direct interaction. Alternatively, as human NINEIN can act as a dynein activator (Redwine et al., 2017), NOCA-2 might act with dynein to recruit γ-tubulin to the MTOC vesicles (Liang et al., 2020). However, we did not observe defects in MTOC vesicle clustering in the *noca-2* mutant as was reported for the dynein mutant (Liang et al., 2020), nor did we see obvious changes in dynein recruitment to the vesicles (Supplemental figure 5C). Therefore, the precise function of NOCA-2 on these MTOC vesicles is still unclear.

In mammals, CAMSAP1, CAMSAP2 and CAMSAP3 all recognize and protect microtubules minus-ends against depolymerization and are important for neuron polarization (Jiang et al., 2014; Pongrakhananon et al., 2018; Yau et al., 2014; Zhou et al., 2020). However, their behavior at the minus-end is different: CAMSAP1 concentrates at the outermost ends and tracks the growing microtubule minus-ends, while CAMSAP2 and CAMSAP3 are stably deposited on the microtubule lattice, forming stretches from the minus-end and stabilizing MT lattices against depolymerization (Hendershott and Vale, 2014; Jiang et al., 2014). In *Drosophila* neurons, Patronin behaves similar to CAMSAP1 that tracks and controls the minus-end growth and helps to populate dendrites with minus-end out microtubules (Feng et al., 2019). In *C. elegans* PLM neurons, microtubule minus-ends growth was also reported in the posterior process (Puri et al., 2021). Here, we found that most of GFP::PTRN-1 (CAMSAP) puncta were highly immobile in mature PVD neurons (Supplemental figure 6D). However, we did observe a small population of anterograde moving PTRN-1 puncta in the shaft of the anterior dendrite (Supplemental figure 6D). The speed of these movements was slower than the typical plus-end growth speeds (Supplemental figure 6E) and may represent microtubule minus-end growth although theses did not overlap with EBP-2::GFP as was the case in *Drosophila* (Supplemental figure 6C)(Feng et al., 2019). However, the PTRN-1 dynamics we observed in the distal segment are much slower and less processive (Figure 4I). This suggest that although minus-end growth may take place in the PVD neuron, we do not expect this to be a major contributor to the dynamics observed in the distal segment.

To organize dendritic microtubules minus-end out, the importance of localizing the MTOC vesicles to the growth dendrite tip is apparent. However, the precise mechanism and the contribution of PTRN-1 (Camsap) to this process is still unclear. One potential model is that PTRN-1 (CAMSAP) stabilizes a specific subset of microtubules that are then used for forward translocation of the MTOC vesicles, e.g. by the UNC-116 (kinesin-1) motor over the short plus-end out microtubules in the tip (Liang et al., 2020). However, we found that PTNR-1 localizes in a punctate pattern surrounding the MTOC vesicles and co-migrate with the vesicles upon dendrite growth. This may suggest that microtubule nucleation is connected to microtubule minus-end stabilization and argues to also consider alternative models where e.g. pushing or pulling on microtubules may translocate the MTOC vesicles forward. Such forces could for example be generated microtubule sliding over other microtubules. In *Drosophila*, kinesin-1 was shown to slide microtubule against other microtubules during early neuronal development, using an extra microtubule binding site in the tail (del Castillo et al., 2015; Lu et al., 2013; Winding et al., 2016b). Although kinesin-1 is also essential in *C. elegans* to organize microtubules minus-end out (Yan et al. 2013), mutating the extra microtubule binding site in using CRISPR did no show microtubule defects arguing against a microtubule-microtubule sliding model for kinesin-1 (results not shown). Alternatively, motors could push or pull the microtubule cytoskeleton forward if anchored to static structures. For example, in axons the dynein motor was shown to push the distal microtubule cytoskeleton forward by anchoring to the cortex and walking to the minus-end of the microtubules (Roossien et al., 2014). Similarly, kinesin-1 may push minus-end out oriented microtubules towards the dendrite tip by walking to the plus-end. Or alternatively, dynein may function at the growing dendrite tip to pull on the short plus-end out microtubules that emanate from the MTOC vesicles, similar to its role to position the centrosomes during cell division (Gusnowski and Srayko, 2011; Kotak et al., 2012; Laan et al., 2012; Nguyen-Ngoc et al., 2007; Schmidt et al., 2017). Interestingly, the forward movements of the MTOC vesicles coincided with longer lived microtubules (Liang et al., 2020), which could represent cortically captured microtubules by dynein. More work is needed to determine the mechanism how the MTOC vesicles are transported anterogradely. Also, how the MTOC vesicles are connected to PTRN-1 stabilized microtubules will be interesting to investigate further. Potentially, NOCA-2 (NINEIN) can connect microtubules to the MTOC vesicles directly as NOCA-1 was reported to bind to microtubules (Wang et al., 2015) as was the *Drosophila* NINEIN homologue, Bsg25D (Kowanda et al., 2016).

Taken together, we propose that the minus-end out microtubule organization in the PVD dendrite follows a two-step model where non-centrosomal microtubules are initially nucleated from MTOC localized γ-tubulin and subsequently stabilized by PTRN-1 (CAMSAP). The minus-end out microtubule population may be further stabilized by cortical anchoring (He et al., 2020) and potentially further amplified by severing proteins such as Katanin and Spastin (Kuo and Howard, 2021). Such a mechanism may also take place in other tissues such as the worm epidermis, where PTRN-1 and NOCA-1 (NINEIN) were found to also act in parallel to organize the non-centrosomal microtubules (Wang et al., 2015). In contrast, in the *Drosophila* fat body, both Patronin (CAMSAP) and NINEIN act independent of γ-Tubulin to assemble non-centrosomal microtubules (Zheng et al., 2020). This indicates that the functional connection between microtubules nucleation and stabilization may vary between cell types and organisms.

## Materials and methods

### C. elegans strains and culturing

Strains were cultured at 15°C or 20°C using OP50 Escherichia coli as a food source and imaged at room temperature. To image early developing PVD neurons, the adult animals were grown at 15°C for at least 48 hr and L2-L3 stage progeny were picked for imaging. The *noca-2(hrt28*) allele used in this study was made using CRISPR-mediated mutagenesis, which deletes the entire NOCA-2 fragment (Supplemental figure 1C). *noca-2(hrt31[GFP]*) was obtained by using CRISPR-based genome editing (Dokshin et al., 2018). The strains: TV21539[tba-1(ok1135) I; wyEx8784[Punc-86::gfp::tba-1; Punc-86: mCherry:: PLCdeltaPH] and *TV25056[wyEx9975[Punc-86::gfp::rab-11.1 cDNA; Punc-86::mCherry::PLCdeltaPH]* were gifts from Dr. Kang Shen (Liang et al., 2020). The strain *gip-1(wow25[tagRFP-t::3xMyc::gip-1]) III* was a gift from Dr. Jessica L. Feldman (Sallee et al., 2018). The strain *ntuIs6[Pdes-2::unc-116(G237A)::mCherry; Pdes-2::unc-104(E250K)::gfp; Podr-1::GFP]III*, was a gift from Dr. Chan-Yen Ou (Chen et al., 2021). The plasmid *Pdes-2::unc-116(G237A)::gfp* was cloned in *Pdes-2::UNC-116(G237A)::mCherry (pCH9*) that was provided by Dr. Chan-Yen Ou (Chen et al., 2021). All trains used in this study are listed in table S1.

### DNA plasmids and gRNA

The DNA plasmids that were used to generate transgenic *C. elegans* strains and the detailed cloning information are listed in table S1. The gRNA used to make *noca-2* mutant and GFP knock in stains are listed in table S1. *Pmyo2::mcherry* (5ng/μl) was used as co-injection marker to generate extrachromosomal strains.

### Microscopy

For all imaging, *C. elegans* were mounted on 5% agarose pads with 10 mM tetramisole or 5 mM Levamisole solution in M9 buffer. All live imaging was performed within 60 min after mounting on a Nikon Eclipse-Ti microscope and with a Plan Apo VC, 60×, 1.40 NA oil or a Plan Apo VC 100 × N.A. 1.40 oil objectives (Nikon). The microscope is equipped with a motorized stage (ASI; PZ-2000), a Perfect Focus System (Nikon), ILas system (Roper Scientific France/PICT-IBiSA, Curie Institute) and uses MetaMorph 7.8.0.0 software (Molecular Devices) to control the camera and all motorized parts. Confocal excitation and detection is achieved using a 100 mW Vortran Stradus 405 nm, 100 mW Cobolt Calypso 491 nm and 100 mW Cobolt Jive 561 nm lasers and a Yokogawa spinning disk confocal scanning unit (CSU-X1-A1N-E; Roper Scientific) equipped with a triple-band dichroic mirror (z405/488/568trans-pc; Chroma) and a filter wheel (CSUX1-FW-06P-01; Roper Scientific) containing BFP (ET-DAPI (49000), GFP (ET-GFP (49002)) and mCherry (ET-mCherry(49008)) emission filters (all Chroma). Confocal images were acquired with a QuantEM:512 SCEMCCD camera (Photometrics) at a final magnification of 110 nm (60x objective), 67 nm (100x objective) per pixel, including the additional 2.0x magnification introduced by an additional lens mounted between scanning unit and camera (Edmund Optics). All EBP-2::GFP and TBA-1::GFP imaging was performed at one frame per second (fps)

For the analysis of PVD neuron morphology, images were acquired using an LSM700 (Zeiss) confocal with a 20x NA 0.8 dry objective using the 488 nm and 555 nm laser lines.

### Quantitative image analysis

Image processing and analysis was done using ImageJ (FIJI) to create kymographs or merged sum intensity images for the intensity profile measurement. Statistical analysis and graphs were made in GraphPad Prism software version 8.0.

To quantify EBP-2::GFP growth orientation, kymographs were made using Kymograph Builder plugin in ImageJ. The retrogradely and the anterogradely growing EBP-2::GFP were manually counted. To quantify the microtubule polarity in the mature PVD dendrite, imaging was performed in the proximal dendrite. For the developing PVD dendrite, the whole anterior PVD region was imaged. To quantify EBP-2::GFP growing speed and frequency (supplemental figure 3A-B) only the distal region (20 μm) was quantified.

To quantify microtubule nucleation (Figure 1G and Supplemental figure 1D), kymographs were made for the whole of anterior dendrite in the developing neurons. When EBP-2 comets mainly grew from the distal anterior dendrites the animal was classified as “distal dendrite nucleation”; when the EBP-2 comets mainly grew from cell body theses were classified as “cell body nucleation” and when EBP-2 comets grew in both directions these were classified as “Mix”.

To quantify the PVD branch complexity, the entire PVD dendrite was divided into 4 segments: one posterior segment (−1) and the anterior segment was divided into three equal length anterior segments (+1, +2 and +3)(Figure 2F). The “branch complexity” index calculation was based on a previous study definition (Taylor et al., 2015b).

To measure the GIP-2 cluster intensity profiles (Figure 3I), cytosolic mKate2 was used to visualize the dendrite tip. We drew a 50 px-wide line from the tip of the anterior dendrite to measure the intensity profile of the GIP-2 cluster and the background intensity was subtracted from the region next to the neuron devoid of gut granule autofluorescence.

## Acknoledgements

We thank Mike Boxem and Sander van den Heuvel (Utrecht University, The Netherlands) for advice, *C. elegans* reagents and infrastructure. We thank Bart de Haan for helping with cloning. We thank Kang Shen for helpful suggestions and sharing of strains. We thank Jessica Feldman, Chan-Yen Ou and Alexander Dammerman for kind sharing of *C. elegans* strains and reagents. Some strains were provided by the CGC, which is funded by the NIH Office of Research Infrastructure Programs (P40 OD010440) and some by the National Biorescource Project. We thank WormBase for curating and making available data related to *C. elegans.* And we thank Amelie Freal for giving feedback on this manuscript.

This work was funded by the Nederlandse Organisatie voor Wetenschappelijk Onderzoek (NWO) (NWO-ALW-VICI 865.10.010 to C.C.H.), by the European Research Council (ERC Consolidator Grant 617050 to C.C.H.) and by the Chinese Scholarship Council (CSC) to L.H.

## Author Contributions

LH and MH designed the project; LH and LvB performed the *C. elegans* experiments, supervised by MH and CH; LH and MH wrote the manuscript.

## Conflict of Interests

The authors declare that they have no conflict of interest.

## Supplemental Figure legend

**Supplemental Figure 1.**
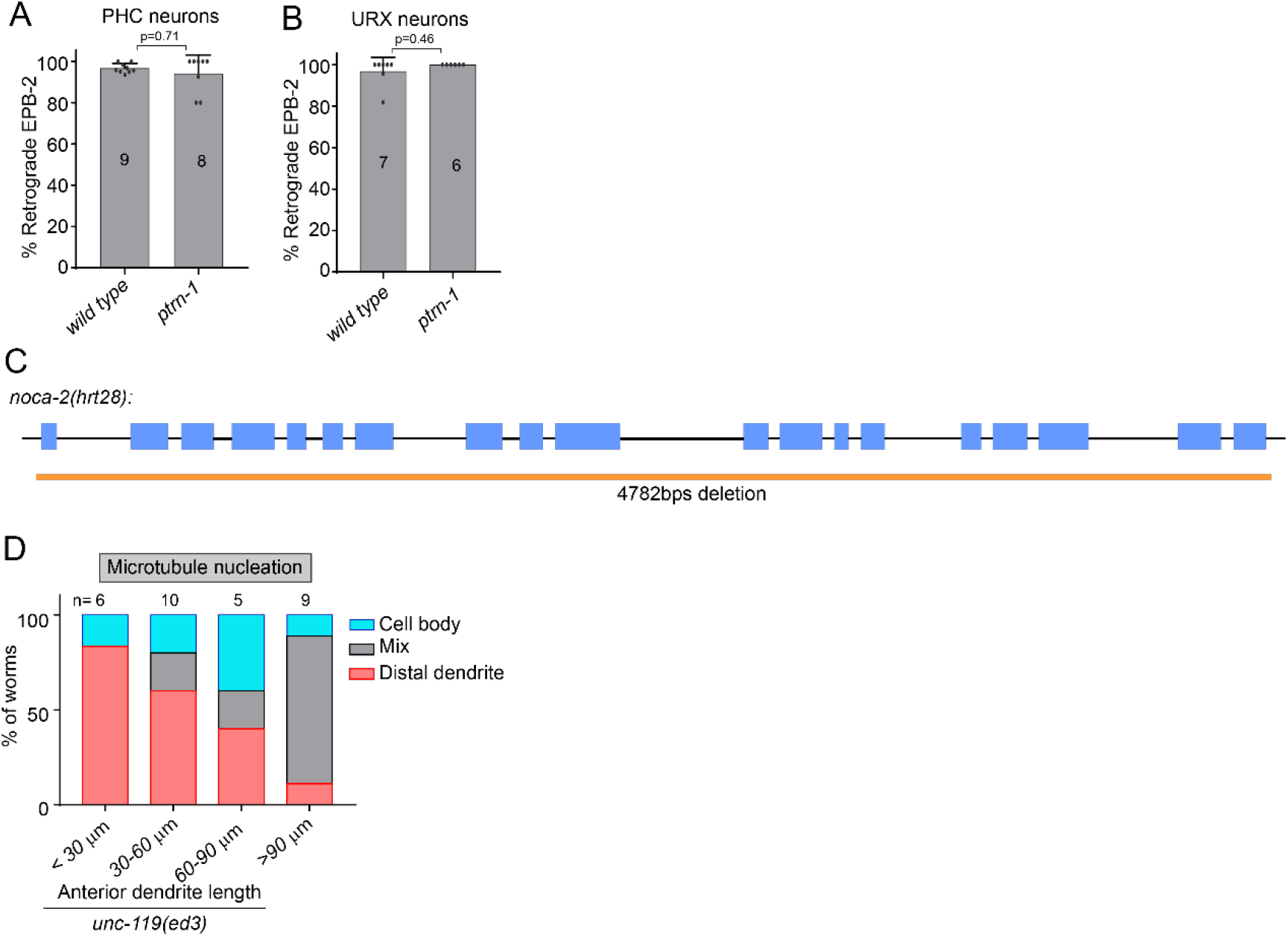
(A-B) Quantification of the percentage of retrograde EBP-2::GFP growth events in the ciliated PHC dendrites (A) and the non-ciliated URX dendrites (B) in wildtype and *ptrn-1* mutant. Error bars represent SD; statistical analysis is followed by unpaired student T-test. Number of analyzed animals is indicated. (C) Gene structure of the *noca-2* gene and *hrt28* deletion. (D) The quantification of microtubule nucleation during neuron developing in the *unc-119* mutant.

**Supplemental Figure 2.**
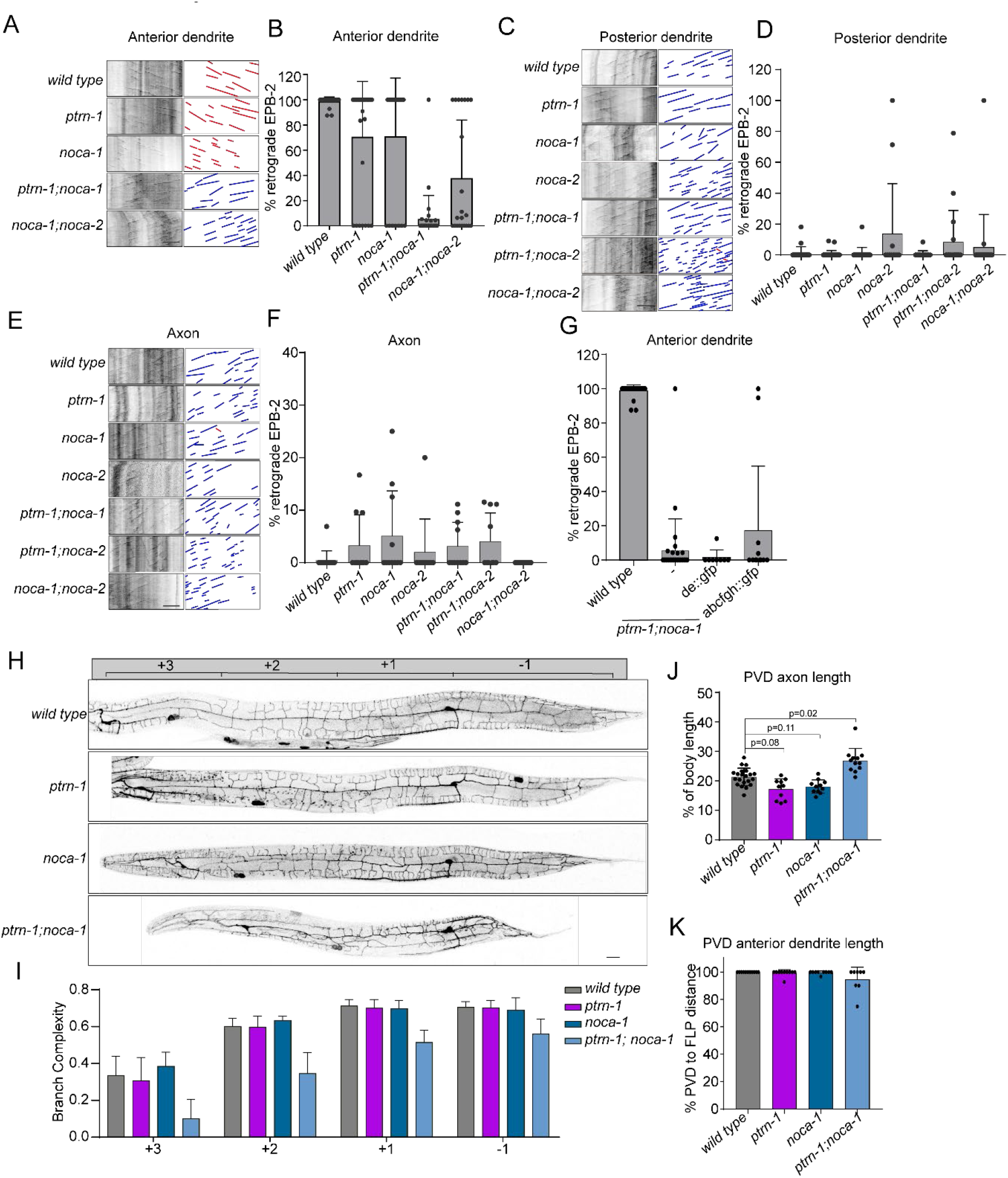
(A-F) Representative kymographs and quantification of microtubule polarity in the mature PVD neurons in the indicated mutants using EBP-2::GFP; The percentage of retrograde growing events in the anterior dendrite (A-B), in the posterior dendrite (C-D) and in the axon (E-F). Scale, 5 μm. (G) Quantification microtubule polarity using EBP-2::GFP in the anterior dendrite of wildtype and *ptrn-1;noca-2* mutants with or without two tagged NOCA-1 rescue constructs(Wang et al., 2015). (H-J) Quantification of the PVD morphology. (H) Representative examples of the PVD morphologies in the indicated mutants. Scale, 20 μm. Quantification of (I) the PVD dendritic branch complexity(Taylor et al., 2015a) (J) the relative axon length in the ventral nerve cord; (K) the relative length of the anterior dendrite. For microtubule polarity analysis, the animals were from L4 to young adult stage. For PVD morphology analysis, only young adult stage animals were analyzed. Error bars represent SD; statistical analysis, Kruskal-Wallis test followed by Dunn’s multiple.

**Supplemental Figure 3.**
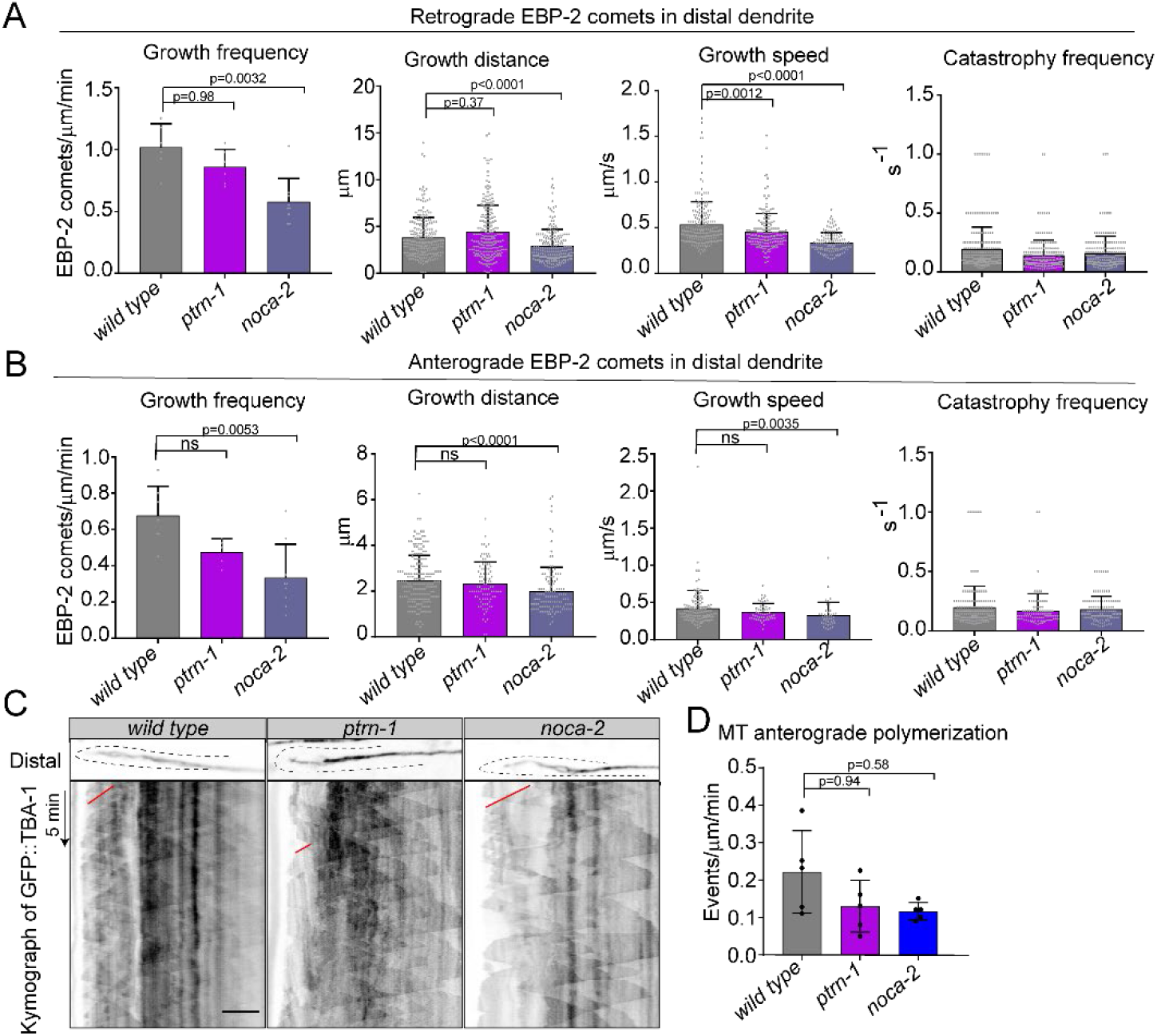
(A-B) Quantification of retrograde (A) and anterograde (B) microtubule plus-end dynamics in the distal 20 μm of the anterior PVD dendrite during neuron development using the plus tip marker EBP-2::GFP. For *ptrn-1* and *noca-2* mutants, only animals that retained distal microtubule nucleation were quantified. For speeds only growth events of >2μm were considered. Scale, 5μm. Error bars represent SD; statistical analysis, Kruskal-Wallis test followed by Dunn’s multiple comparisons test. Number of analyzed animals is indicated. (C) Representative kymographs of GFP::TBA-1 in the distal anterior dendrite. Red lines, growing plus-end out MTs in distal region. scale bar, 5μm. The distal anterior PVD dendrites are indicated with dashed lines. (D) Quantification of plus-end out microtubule polymerization frequencies in the distal region of the growing anterior dendrite. Only animals that have distal microtubule nucleation were considered. Error bars represent SD; statistical analysis, Kruskal-Wallis test followed by Dunn’s multiple comparisons test. 5 animals for each group were analyzed.

**Supplemental Figure 4.**
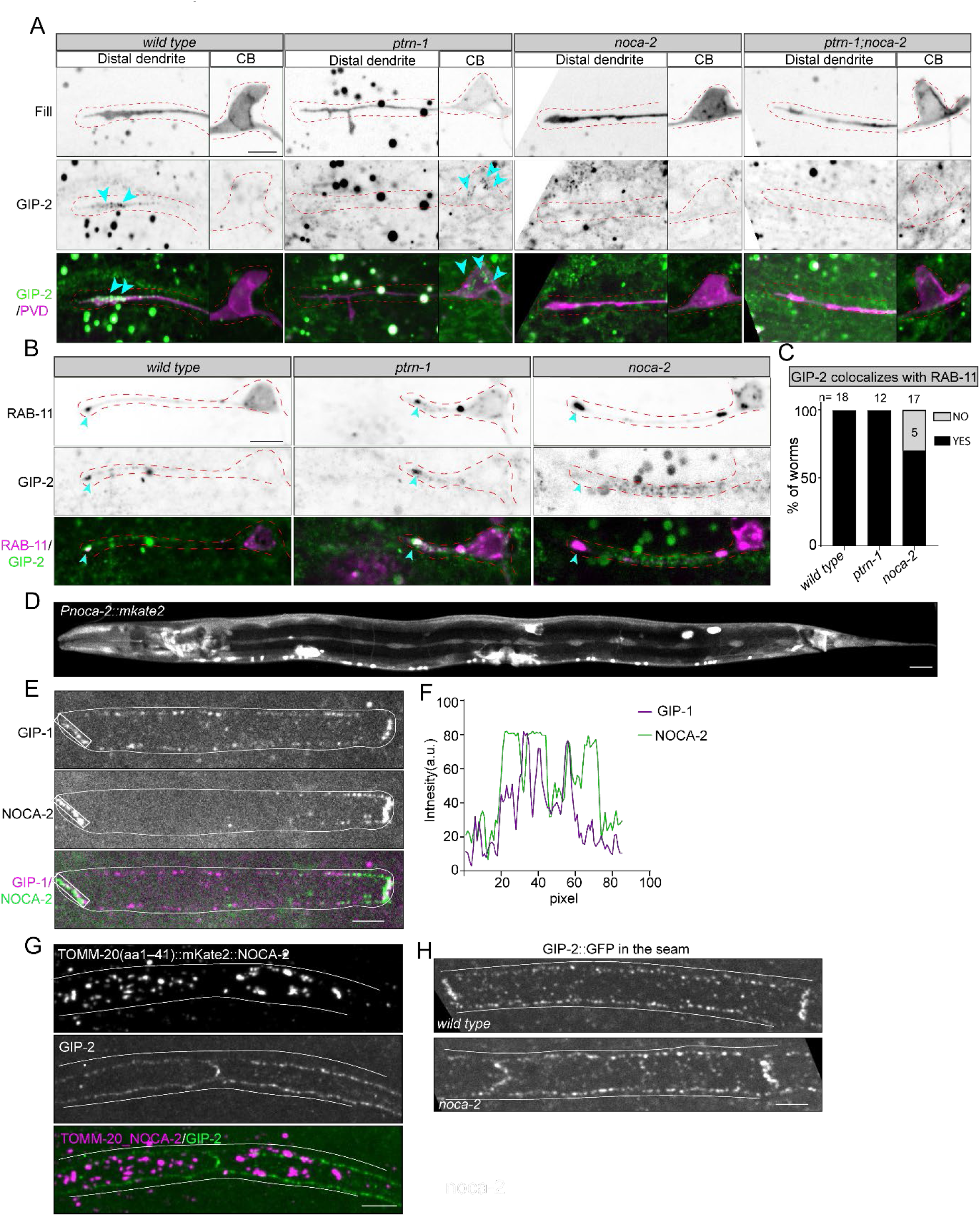
(A) Representative examples of endogenously tagged GIP-2::GFP in the PVD neuron. GIP-2 accumulated in the cell body in the *ptrn-1* mutant (second panel) and without obvious GIP-2 accumulation in *noca-2* mutant (third panels) and *ptrn-1;noca-2* mutants (last panels). Green: GIP-2, magenta: PVD neuron fill. GIP-2 puncta are indicated with blue arrowheads. Scale, 5 μm. The developing neurons are indicated with red dashed lines. (B) Example images of mKate2::RAB-11 (magenta) and GIP-2::GFP (green) co-localization in the distal segment of the growing PVD anterior dendrite. The localization of RAB-11 and/or GIP-2 in developing anterior dendrite is indicated with arrowheads. Scale, 5 μm. The PVD neuron is indicated with red dashed lines. (C) Quantification of the number of animals in which mKate-2::RAB-11 colocalizes with GIP-2::GFP in the distal segment of the growing PVD anterior dendrite; gray: the percentage of animals that have RAB-11 accumulated in distal dendrites but without GIP-2 accumulation. Number of analyzed animals is indicated. (D) The expression of mKate-2 driven by 2kb of *noca-2* promoter sequence. Scale, 20 μm. (E-F) Examples of localization(E) and the intensity quantification(F) of endogenous NOCA-2 and GIP-1 in epidermal seam cells. Scale, 5 μm. The epidermal seam cells are marked with white lines. (G) Representative example images of the GIP-2(green) localization upon artificial NOCA-2 (Magenta) re-localization to mitochondria by fusing it to TOMM-20(1-41 amino acids). Scale, 5 μm. The epidermal seam cells are marked with white lines. (H) The localization of GIP-2 in epidermal seam cells in wild type (upper panels) and in *noca-2* mutant (lower panels). Scale, 5 μm. The epidermal seam cells are marked with white lines.

**Supplemental Figure 5.**
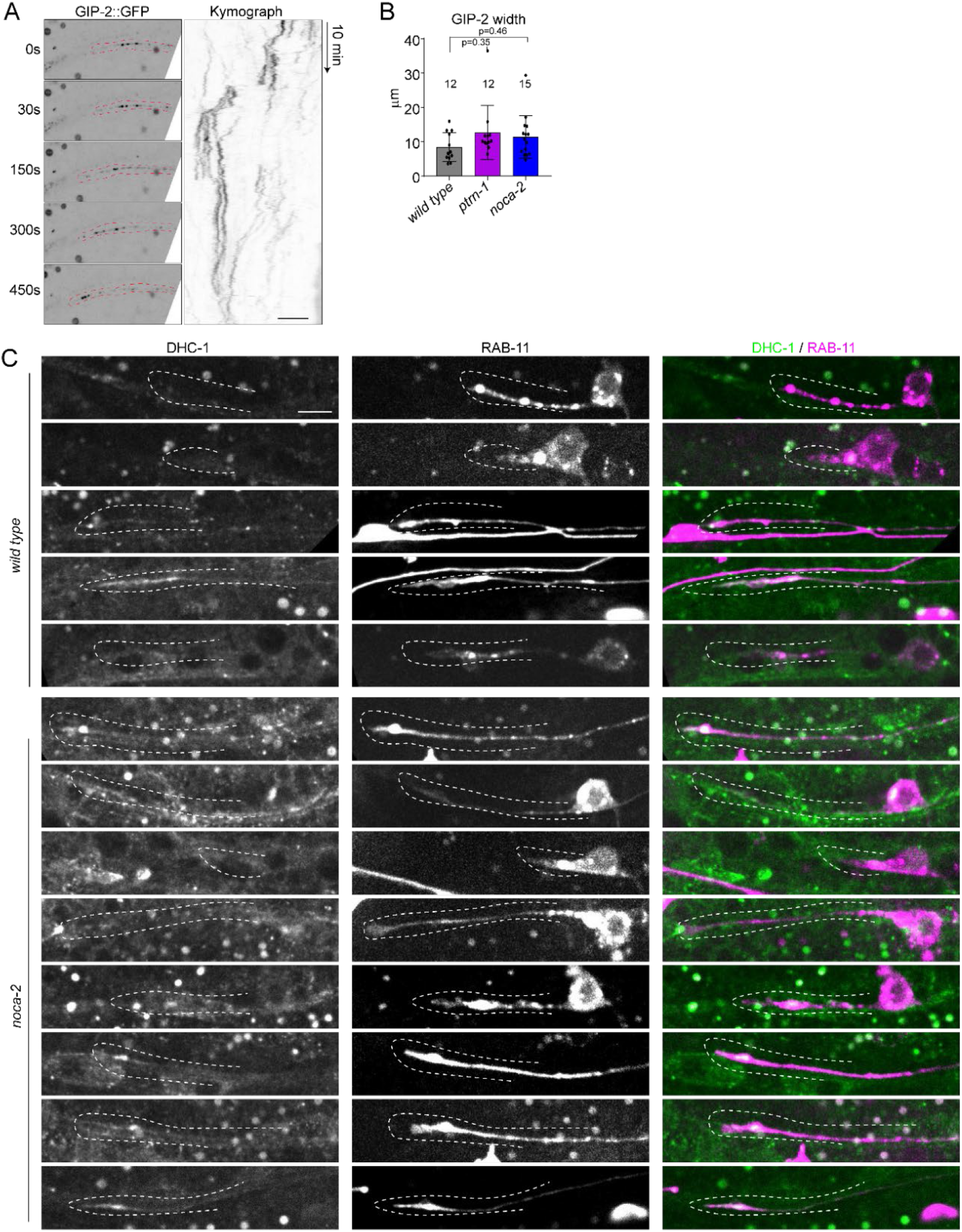
(A) Stills and kymograph of endogenous GIP-2::GFP localization over time (left panel) in growing PVD anterior dendrite. Scale, 5 μm. The distal dendrite is indicated with red dashed lines. (B) Quantification of GIP-2 cluster width in the growing PVD anterior dendrite. (C) Multiple examples of endogenous DHC-1::GFP (green) and PVD expressed RAB-11::mKate2 (magenta) in the growing PVD anterior dendrite. Scale, 5 μm. The distal anterior PVD dendrites are indicated with white dashed lines.

**Supplemental Figure 6.**
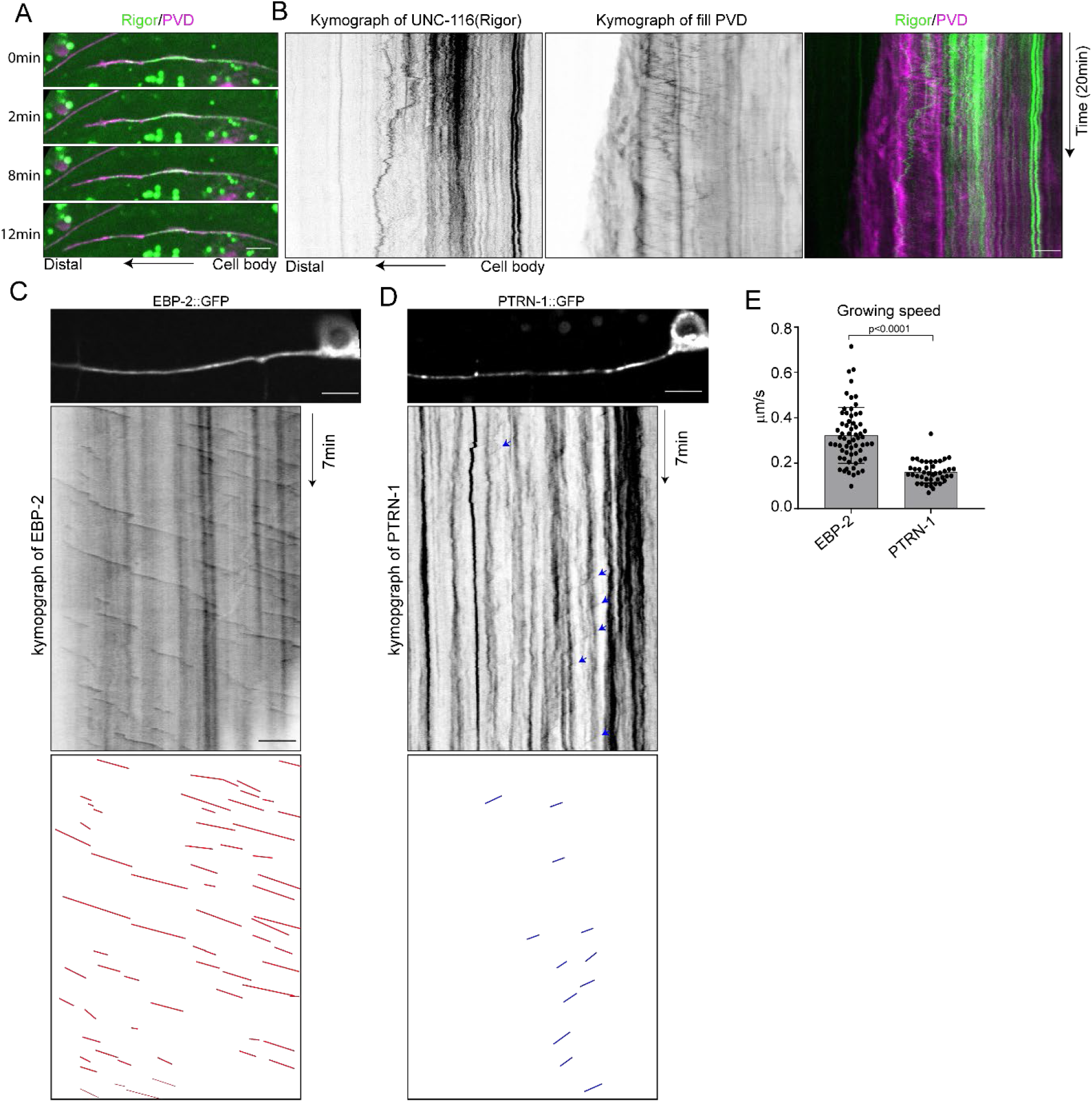
(A-B)) Stills (A) and kymographs (B) of UNC-116(rigor)::GFP localization in the PVD anterior dendrites (A, green). Myristoylated mKate2 was used as a fill (magenta). Scale, 5 μm. (C-D) Representative example kymograph of EBP-2::GFP (C) and mKate2::PTRN-1 (D) in the mature PVD anterior dendrite. Examples of moving PTRN-1 puncta are indicated with blue arrowheads. Scale, 5 μm. (E) Quantification of EBP-2::GFP growth speed and PTRN-1 moving speed in the mature PVD anterior dendrite. Error bars represent SD; statistical analysis is followed by unpaired student T-test.

## Videos

Video 1: EBP-2::GFP comets in the developing anterior PVD dendrite in wild type and indicated mutants. Time, min:sec

Video 2: GFP::RAB-11 (green) cluster movement in growing PVD anterior dendrite (magenta) in wild type and indicated mutants. Time, min:sec

Video 3: The accumulation and dynamic of NOCA-2::GFP in distal developing anterior dendrites. Time, min:sec

Video 4: Co-movement of PTRN-1::mKate2 (magenta) and the MTOC marker (GIP-2::GFP, green) in developing PVD anterior dendrite. Time, min:sec

## Notes

### Competing Interest Statement

The authors have declared no competing interest.

